# High-fidelity CRISPR genome editing of single-nucleotide mutation with near-complementary guide RNA via enhanced target binding kinetics

**DOI:** 10.1101/2025.09.02.671676

**Authors:** Hyomin K Lee, Seohyun Kim, Hyeon Jong Yu, Juyoung Hong, Taegeun Bae, Yohan An, Chang Ho Sohn, Woo Chang Hwang, Chul-Kee Park, Seung Hwan Lee, Hye Ran Koh, Junho K Hur

**Author notes:** **Co-responding authors** Chul Kee Park, Seung Hwan Lee, Hye Ran Koh, Junho Hur. These authors contributed equally to this work (Hyomin K Lee, Seohyun Kim).

## Abstract

The CRISPR-Cas9 system is a powerful genome editing tool capable of precisely recognizing and cleaving specific DNA sequences, and has been extensively investigated as a strategy for correcting mutations associated with genetic diseases and cancer. However, conventional CRISPR genome engineering often fail to discriminate single-nucleotide mutations from wild-type alleles when the mutation is located outside the protospacer adjacent motif (PAM) sequence. To address this limitation, we developed a RNA engineering approach for designing near-complementary single guide RNA (sgRNA) that contain intentional mismatches within the seed region of the sgRNA. Single molecule kinetic analyses showed that the near-complementary sgRNA selectively reduces the binding affinity of CRISPR ribonucleoprotein complex by via differentiated increment in the dissociation rates to the wild-type target DNA compared to the mutant allele. The engineered kinetic characteristics of near-complementary sgRNAs enable highly specific genome editing of single-base mutations without reliance on PAM proximity. We demonstrate the application of the strategy to the a cancer-specific single-nucleotide G228A (-124C > T) mutation in the *TERT* promoter, frequently found in glioblastomas and other tumors, that does not generate a canonical PAM sequence. Our near-complementary sgRNA successfully induced selective editing of the mutant allele while sparing the wild-type sequence. Furthermore, single-molecule fluorescence resonance energy transfer (smFRET) analyss revealed distinct differences in binding kinetics between mutant and wild-type DNA, providing kinetic insight into the discrimination process. We conclude that the near-complementary sgRNA CRISPR editing strategy facilitates precise PAM-independent targeting of single-nucleotide mutations without protein engineering and offers a molecular framework for expanding the specificity and applicability of CRISPR-based genome and epigenome editing technologies.

## Background

Single-nucleotide mutations represent the most prevalent class of pathogenic genetic alterations and are a major cause of hereditary disorders as well as oncogenic driver events in cancers such as glioblastoma and melanoma[1-6]. The CRISPR–Cas9 system has emerged as a genome editing tool owing to its programmability and precision in targeting DNA sequences[7, 8]. Cas9 identifies genomic targets through base pairing between a single guide RNA (sgRNA) and a complementary 20-nucleotide sequence located adjacent to a protospacer-adjacent motif (PAM), typically NGG for *Streptococcus pyogenes* Cas9 (SpCas9)[7, 9-11].

Although Cas9 recognizes the target sequence through PAM dependence and guide RNA complementarity, perfect allele discrimination is often not achieved in practice. In previous studies, it has been demonstrated that a single mismatch between the target sequence and the sgRNA can lead to unintended cleavage of both mutant and wild-type alleles[12-14]. In some instances, CRISPR–Cas9 can distinguish between mutant and wild-type alleles when single-nucleotide mutations generate a de novo PAM site, thereby enabling allele-specific editing[15-17]. However, such cases are relatively rare, as the majority of disease-associated single-nucleotide mutations do not derive PAM motif. Furthermore, the inherent mismatch tolerance of Cas9—especially at PAM-distal positions—limits its ability to discriminate between single-base differences, often resulting in off-target editing of wild-type alleles[12, 18]. To address this challenge, previous studies have explored sgRNA sequence engineering to enhance mismatch sensitivity and reduce editing activity at wild-type loci[19, 20]. While these approaches have shown some promise, the mechanistic basis of their improved specificity remain poorly understood, limiting their applicability to PAM-independent CRISPR targeting of the single-base mutations.

In this study, we present a sgRNA rational design strategy that enables precise PAM-independent genome editing with highly effective discrimination of single-nucleotide mutations. Using the clinically relevant G228A (−124C>T) mutation in the *TERT* promoter—a cancer-specific substitution that does not produce a canonical PAM—we generated near-complementary sgRNAs By introducing intentional mismatches within the seed region of the sgRNA [21]. Ribonucleoprotein complexes of CRISPR-spCas9 and near-complementary sgRNAs were capable of selectively targeting mutant alleles while avoiding cleavage of the wild-type sequence. Single-molecule Förster resonance energy transfer (smFRET) analyses further revealed that discrimination arises from differences in binding stability and dwell time, offering kinetic insight into the molecular mechanism underlying single-base resolution editing[9, 22]. The results show that genome editing by near-complementary sgRNA design achieves high allele specificity in both in vitro and in human cells. As the RNA-engineering framework is applicable for allele-specific CRISPR tools without protein engineering, it may facilitate the development of CRISPR gene therapies to address dominant-negative or gain-of-function pathogenic mutations.

## Results

Previous clinical studies have demonstrated a high prevalence of the monoallelic *TERT* G228A mutation across various cancer cell lines [23, 24]. This study aimed to evaluate the specificity of CRISPR-Cas9 in targeting the single-base *TERT* G228A substitution. We analyzed sgRNA sequence complementarity to the *TERT* G228A mutation versus the wild-type sequence (Additional file 1: Fig. S1A). Using sgRNA-A, which is designed to target the *TERT* G228A mutation, we performed genome editing in Huh7 (*TERT* G228A) and HEK293T (wild-type) cell lines, following transfection with Cas9-encoding plasmids. Mutation rates were quantified using targeted deep sequencing (Additional file 1: Fig. S1B). Surprisingly, CRISPR-Cas9 directed by sgRNA-A resulted in significant insertions and deletions (indels) in both Huh7 and HEK293T cells, indicating significant off-target genome editing activity in cells lacking the G228A mutation. Similarly, sgRNA-G, complementary to the wild-type sequence, caused notable genome editing in both cell lines suggesting inadequate discrimination between the mutated and wild-type sequences.

Next we asked if better distinction of the TERT G228A single-base difference could be achieve by employing the eSpCas9 high-fidelity CRISPR system[20]. To assess the specificity of eSpCas9 to target *TERT* G228A mutation, sgRNAs were co-transfected with corresponding reporters (Additional file 1: Fig. S1C)[25, 26]. Both sgRNA-G and sgRNA-A produced similar mRFP–EGFP double-positive cell ratios of 38.4% and 42.4% with the *TERT* G228A reporter, and 39.3% and 41.8% with the wild-type reporter, respectively. These findings implied that genome editing by eSpCas9 did not show significant target selectivity of *TERT* G228A single-base mutation.

We posited that introducing an additional mismatch in the seed region of sgRNAs targeting the *TERT* G228A mutation—resulting in two mismatches relative to the wild-type sequence— could enhance the specificity of CRISPR-Cas9 genome editing (Fig. 1A). To this end, we engineered a series of sgRNAs (sgRNA-A1 to sgRNA-A6), each incorporating a deliberate mismatch in the seed region of the original sgRNA-A (Additional file 2: Table S1). Subsequent in vitro cleavage assays employed these sgRNAs alongside the Cas9 protein to treat PCR amplicons derived from the genomic DNA of *TERT* G228A-mutant (Huh7 and KMH-2) and wild-type (HEK293T and HeLa) cells (Additional file 1: Fig. S2A). In our assays, sgRNA-A1 emerged as the most specific guide RNA, efficiently targeting the *TERT* G228A-mutated sequence while demonstrating no detectable cleavage activity on the wild-type sequence, consisted with our hypothesis of its high specificity (Fig. 1B). In contrast, sgRNA-A, A4, and A5, while also targeting the *TERT* G228A mutation, showed varying degrees of cleavage in wild-type sequences, indicative of reduced targeting precision (Additional file 1: Fig. S2B). As sgRNA-A1 showed superior discrimination, we selected the sgRNA to conducted the subsequent experiments.

**Fig 1.**
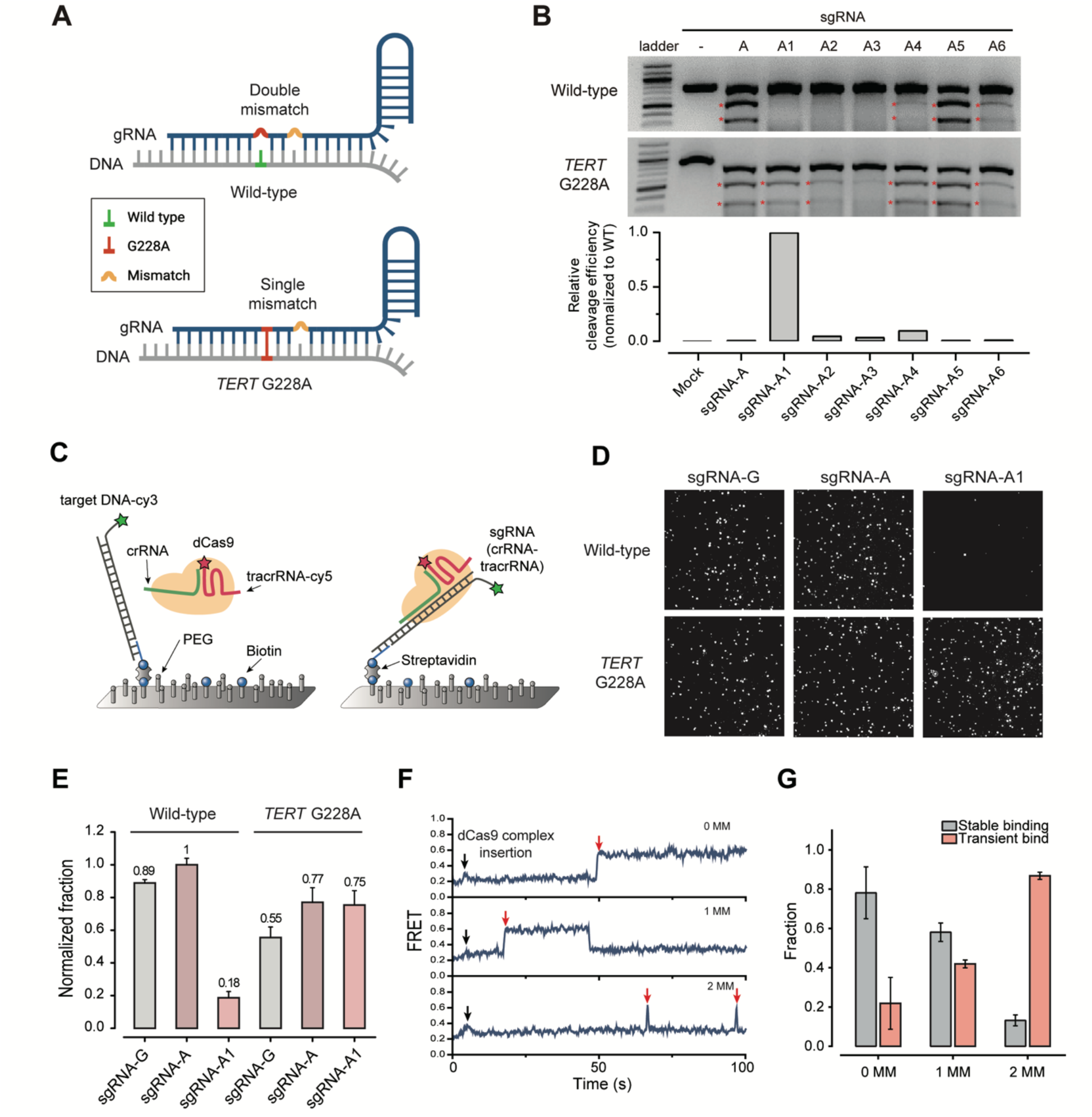
Near-complementary sgRNAs increases the specificity of CRISPR ribonucleoprotein complex to single-nucleotide mutations by enhancing the binding dynamics. **(A)** Schematic of near-complementary sgRNA targeting *TERT* G228A mutation. The green line indicates wild-type DNA (top) and the red line indicates *TERT* G228A mutant DNA (bottom). The yellow bulge shows the mismatches. **(B)** *In vitro* cleavage assay using SpCas9.Lane (1), Mock (SpCas9 only); lane (2), SpCas9 with sgRNA-A; lanes (3) to (8), SpCas9 with near-complementary gRNA-A1 to -A6. Cleaved DNA fragments are indicated with red asterisks. The accompanying bar graph shows the relative cleavage efficiencies for *TERT* G228A mutant versus wild-type templates.Cleavage ratios were quantified from band intensities and normalized to the wild-type cleavage efficiency for each sgRNA. **(C)** Schematic of the single-molecule binding assay to detect the binding of dCas9 complex with *TERT* DNA using total internal reflection fluorescent (TIRF) microscopy. Fluorescence resonance energy transfer (FRET) appeared upon the binding of the Cy5-labeled dCas9 complex with the Cy3-labeled *TERT* DNA, which was immobilized onto the PEG-coated surface. **(D)** Representative single-molecule fluorescence images showing the immobilized Cy5-labeled dCas9 complex with three different crRNAs that bound to *TERT* DNA. **(E)** Normalized fraction of dCas9 complex bound to *TERT* DNA. The binding of the dCas9 complex to *TERT* DNA decreased significantly only for the double DNA–crRNA mismatches (wild-type DNA and crRNA-A1). **(F)** Representative FRET time traces showing the binding of the dCas9 complex to *TERT* DNA (red arrows) for 0, 1, and 2 DNA–crRNA mismatches (0 MM, 1 MM, and 2 MM, respectively) upon the addition of the dCas9 complex (black arrows). **(G)** The fraction of stable and transient binding according to the number of DNA–crRNA mismatches. The binding events with a dwell time longer than 1.5 s were considered for stable binding.

We next sought to assess biochemical mechanism of the significantly reduced cleavage of wild-type DNA by the Cas9-sgRNA-A1 complex compared with the *TERT* G228A mutant. To this end, we employed a single-molecule fluorescence binding assay using catalytically inactive Cas9 (dCas9). In this assay, the CRISPR ribonucleoprotein complexes were assembled from fluorescently labeled dCas9 protein and two RNAs, crRNA and tracrRNA, that anneal together to form a single RNA structure that substitutes for sgRNA. To accommodate the RNA sequence alterations, we utilized several types of crRNAs contained varying mismatches and a common Cy5-labeled tracrRNA. We immobilized Cy3-labeled target DNA, either wild-type or *TERT* G228A mutant, on a PEG-coated surface via biotin– neutravidin linkage. The dCas9 complex, comprising dCas9, crRNA, and Cy5-labeled tracrRNA, was then introduced onto the surface (Fig. 1C). Upon binding, two distinct FRET efficiency peaks were detected at ∼0.30 and ∼0.55 with 532 nm laser excitation, corresponding to two conformational states of the dCas9-DNA complex (Additional file 1: Fig. S3A and S4). Consistent with previous studies, single-molecule FRET trajectories revealed transition between these states, likely reflecting different binding modes related with R-loop formation dynamics (Additional file 1: Fig. S3) [27, 28].

We further quantified the binding affinities of dCas9 complexes for wild-type and mutant *TERT* DNA targets, using crRNAs with varying degrees of complementarity to each target. The dCas9 complexes with perfect matches (crRNA-G with wild-type DNA and crRNA-A with mutant DNA) or a single mismatch (crRNA-G with mutant DNA, crRNA-A with wild-type DNA, and crRNA-A1 with mutant DNA) presented significantly higher density of Cy5 fluorescence spots than those with two mismatches (crRNA-A1 with wild-type DNA) (Fig. 1D and additional file 1: Fig. S4C). Binding affinity was quantified by normalizing the number of Cy5-labeled dCas9 spots to the number ofCy3-labeled DNA spots for each target. This analysis revealed the lowest bound fraction for the double-mismatch condition (crRNA-A1 with wild-type DNA) (Fig. 1E), demonstrating that the presence of two mismatches markedly reduces the binding affinity of dCas9 complex for target DNA, consistent with the observed inefficiency in cleavage.

To investigate the molecular basis for the reduced binding affinity observed with double mismatches, we monitored dCas9-target DNA interactions in real time using single-molecule FRET. FRET signals appeared upon binding of Cy5-labeled dCas9 to Cy3-labeled target DNA and disappeared upon dissociation, allowing direct observation of individual binding and release events (Fig. 1F). Dwell-time analysis for complexes with zero, one, or two mismatches revealed progressively shorter binding durations as the number of mismatches increased, with corresponding dissociation rate of 0.028 s^-1^(0 mismatches), 0.038 s^-1^(1 mismatch), and 0.27 s^-1^(2 mismatches). (Fig. 1F and Additional file 1: Fig. S5). All conditions exhibited both transient and stable binding events, where stable binding was defined as events lasting at least twice the average transient dwell time (Fig. 1G). While most zero-mismatch interactions resulted in stable binding, mismatch introduction markedly reduced this stability (Fig. 1G), implicating that the decreased binding affinity in the double-mismatch dCas9 complex arises from a substantially faster dissociation rate. Together, these findings demonstrate that double mismatches at specific positions markedly reduce the binding affinity compared to a single mismatch and consequently the cleavage activity of the Cas9-sgRNA complex, a property that can be exploited for sensitive mutation detection assays.

To assess the specificity of sgRNA-A1 within a cellular level, we employed a dual fluorescence reporter system that expresses mRFP and EGFP (Fig. 2A). This system incorporates coding sequences for mRFP and EGFP, which are separated by either *TERT* G228A or wild-type target sequences (Additional file 1: Fig. S6A). While mRFP is constitutively expressed in cells, EGFP expression is contingent upon successful genome editing. To evaluate the specificities of sgRNA-G, sgRNA-A, and sgRNA-A1, we co-transfected these sgRNAs alongside the wild-type and G228A reporters. In cells containing the *TERT* G228A reporter, baseline mRFP–EGFP double-positive cells were under 1% of the total cell population. However, transfection with sgRNA-G, sgRNA-A, and sgRNA-A1 yielded double-positive cell ratios of 47.9%, 57.4%, and 40.7%, respectively, indicating comparable genome editing efficiencies among the sgRNAs (Fig. 2B).

**Fig 2.**
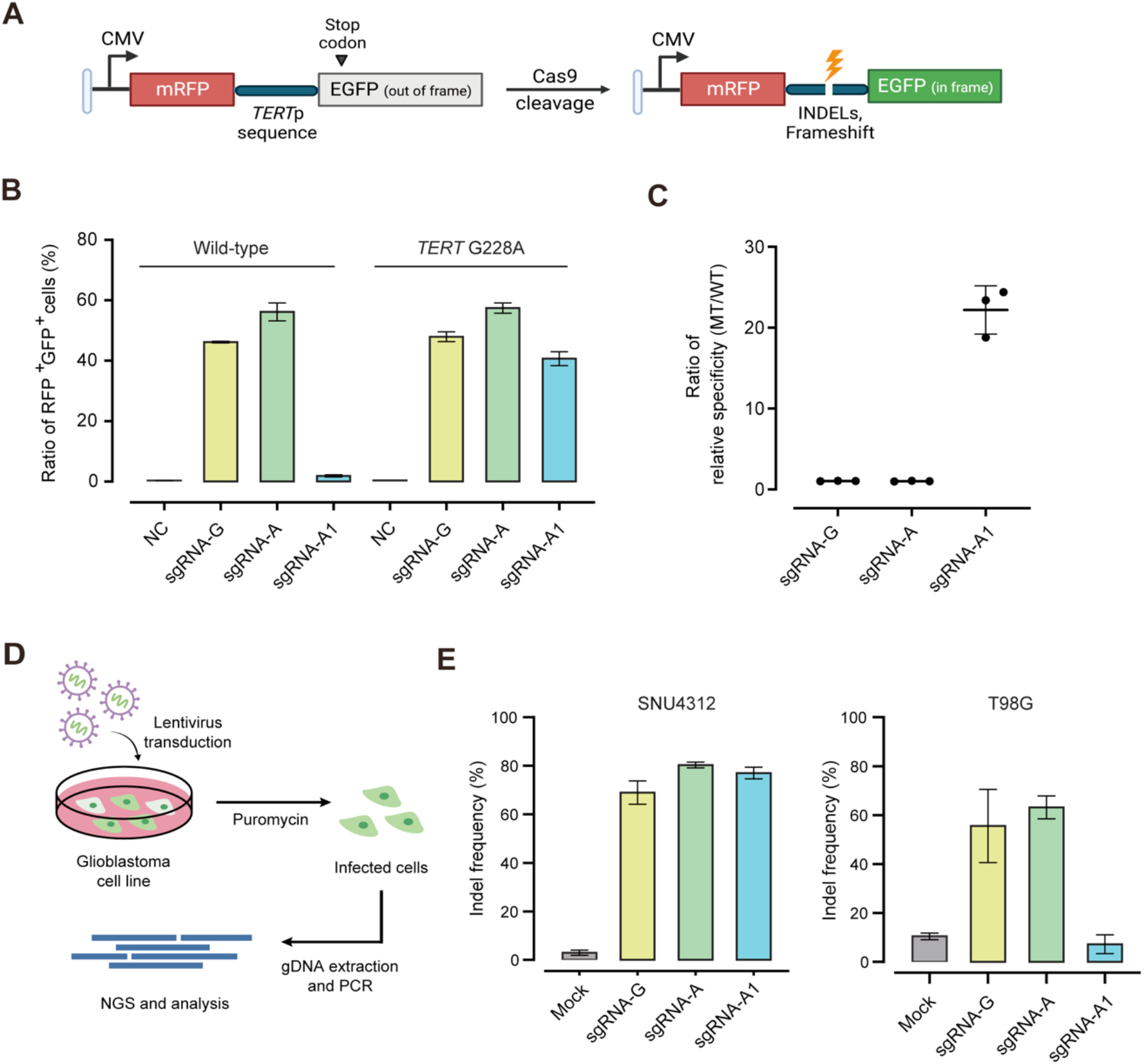
Precision gene editing of single-nucleotide *TERT* G228A mutation by near-complementary sgRNA in fluorescence reporter and endogenous mutant allele in glioblastoma cell lines. **(A)** Schematic of the mRFP–EGFP dual fluorescence reporter system. The target sequences were encoded between mRFP and EGFP. The dual fluorescence reporter had a stop codon in front of the EGFP-encoding sequence. In the case of insertions and deletions (indels) at target sites, the stop codon was frame-shifted, leading to EGFP expression. **(B)** Graphs representing the ratios of the mRFP- and EGFP-expressing populations divided by the entire populations upon wild-type and *TERT* G228A sequence targeting **(C)** Relative specificities, calculated by dividing the indel frequency of the wild-type sequence by that of *TERT* G228A mutant allele **(D)** Schematic of steps of gene editing in endogenous *TERT* G228A mutation in glioblastoma cell line. Genomic DNA was extracted at 20 days post-transduction and selected with puromycin. Mock indicates the background level of insertion and deletions (indels) measured in non-edited cells. Genomic DNA obtained from four independent experiments. **(E)** Comparison of sgRNA target specificity in SNU4312 (G228A) and T98G (wild-type) cells. Targeted deep-sequencing analysis of indel formation at the *TERT* G228A mutation site is shown.

Conversely, the wild-type reporter demonstrated average double-positive ratios of 46.2% and 56.2% when edited by sgRNA-G and sgRNA-A, respectively. Remarkably, sgRNA-A1 exhibited a markedly lower ratio of 1.9%. The specificity of sgRNA-A1 for the G228A mutation over the wild-type sequence was 22.2-fold greater, while sgRNA-G and sgRNA-A showed negligible increases in specificity at 1.04- and 1.02-fold, respectively (Fig. 2C). These results suggested that sgRNA-A1 could distinguish single-base differences between wild-type and the *TERT* G228A mutation with high efficiency, consistent with results of the in vitro cleavage assay.

In exploring the cellular application of near-complementary sgRNA-A1, we aimed to confirm its ability to facilitate specific gene editing. We evaluated the gene-editing efficiency of sgRNA-A1, which is specific to the *TERT* G228A mutation, in a human glioblastoma multiforme (GBM) cell line[29, 30]. The DNA cleavage efficiency of sgRNA-A1 was assessed within this context. We utilized a lentiviral vector encoding CRISPR-Cas9 and the sgRNA-A1 targeting the *TERT* G228A mutation. Lentiviral particles were introduced into SNU4312 and U373 (harboring *TERT* G228A) cell lines, with T98G (containing *TERT* G250A) serving as a negative control (Fig. 2D). Our findings revealed a marked reduction in indel frequencies in T98G cells when edited with sgRNA-A1. In contrast, sgRNA-A1 yielded high indel frequencies in SNU4312 and U373 cells, comparable to the mutant-complementary sgRNA-A (Fig. 2E and Additional file 1: Fig. S7B). This demonstrates that intracellular application of sgRNA-A1 significantly enhances specificity for the *TERT* G228A target. However, sgRNA-A exhibited high editing efficiency across all cell lines tested, irrespective of the presence of the *TERT* G228A mutation, indicating non-specific binding.

To determine the potential for unintended off-target effects by sgRNA-A1, we first conducted a comprehensive in silico human genome-wide search, comparing potential off-target sequences related to sgRNA-A and sgRNA-A1 (Additional file 2: Table. S2). Computational analyses revealed a reduction in potential off-target sites from six to one for sgRNA-A1, excluding matches to wild-type sequences. We further validated the off-target effects in sgRNA-A-and sgRNA-A1-transduced cell lines (SNU4312, U373) using targeted deep sequencing (Additional file 1: Fig. S7C). Administration of sgRNA-A led to approximately 8% indels at off-target sites 2 and 3. In contrast, sgRNA-A1 was associated with less than 0.1% indels at these sites, akin to mock-treated conditions. Notably, off-target site 3 is situated within the coding region of the plexin B1 (PLXNB1) gene, implicated in cell shape regulation and the semaphorin-plexin signaling pathway[31, 32]. However, sgRNA-A1 maintained indel frequencies below 0.1% at all off-target sites, except site 7, which is not proximal to any gene, thus suggesting no significant genomic impact. These results support the conclusion that sgRNA-A1-driven CRISPR gene editing minimizes critical off-target effects in the genome. By applying this method to other genes or targets, it will be possible to easily obtain CRISPR technology with high and elaborate specificity. Based on the results, we conclude that this strategy will be widely applicable to the development of CRISPR gene therapy for diseases, including cancer and genetic disorders, with single-point mutations.

## Conclusion

In this study, we developed and validated a novel PAM-independent genome editing strategy to selectively target single-nucleotide mutations by engineering near-complementary sgRNAs with deliberate seed region mismatches. Using the clinically relevant *TERT* promoter G228A mutation as a model, we demonstrated that this design confers high specificity for the mutant allele while minimizing off-target editing of the wild-type sequence. sgRNA-A1, incorporating a rationally placed mismatch, showed minimal activity on wild-type DNA in both in vitro cleavage, single molecule analyses and cellular reporter assays, while maintaining efficient editing in G228A-harboring cancer cell lines.

Single-molecule FRET analysis revealed that enhanced specificity stems from impaired binding stability and reduced dwell time of the Cas9 complex when mismatched to the wild-type target, providing mechanistic insight into how mismatch tuning in sgRNA can exploit biophysical dynamics for single-base discrimination. Furthermore, genome-wide off-target profiling confirmed a drastic reduction in unintended edits with sgRNA-A1 compared to conventional sgRNAs.

Collectively, our results establish a broadly applicable framework for designing highly specific CRISPR-Cas9 tools capable of discriminating single-nucleotide mutations even in the absence of PAM sequences. We anticipate that this approach has strong potential for safe and precise genome and epigenome editing methods and could be further utilized for developing gene therapies targeting dominant-negative or gain-of-function mutations in cancer and genetic diseases.

## Materials and Methods

### Cell subculture and transfection

HEK293T cells (ATCC) were cultured in DMEM media (Welgene) supplemented with 10% fetal bovine serum (FBS) and 1% antibiotics. Glioma cell lines T98G and U373, sourced from the Korean Cell Line Bank (KCLB), were maintained in RPMI 1640 media (Welgene) containing 10% FBS and 1% antibiotics. SNU4312, glioma cell lines, also from KCLB, were cultivated in Opti-MEM (Gibco) with 10% FBS and 1% antibiotics. For the RFP–GFP dual reporter assay, 1.5 × 10^5 HEK293T cells were seeded in 24-well plates and transfected with 200 ng of SpCas9 plasmid, 200 ng of sgRNA plasmid, and 100 ng of reporter vector using Lipofectamine 2000 (Invitrogen, Waltham, MA, USA). Cells were cultured for 48 hours before FACS analysis for mRFP and EGFP expression.

### In vitro transcription of sgRNA

Target sgRNA sequences (Additional file 2 : Table S1) were synthesized (macrogen) and annealed to create DNA templates for in vitro transcription with T7 RNA polymerase (NEB), alongside a reaction mixture of 10× RNA polymerase reaction buffer, 50 mM MgCl2, 100 mM rNTP, RNase inhibitor (murine), 100 mM DTT, and nuclease-free water. After 6 hours, incubation at 37°C, DNA templates were removed by DNase1 treatment, and sgRNAs were purified using an RNA purification kit (RBC).

### In vitro DNA cleavage assay and calculation of the DNA cleavage efficiency

On- and off-target PCR amplicons were generated from genomic DNA using specific primer sets (Additional file 2 : Table S4). The PCR Amplicons were incubated with purified Cas9 protein and sgRNAs at 37°C for 1 hour in NEB3.1 cleavage buffer. Reactions were halted using stop buffer (100 mM EDTA, 1.2% SDS), and DNA cleavage was visualized on a 1.5% agarose gel. Cleavage efficiencies were quantified 0using ImageJ software based on the formula (intensity of the cleaved fragment / total fragment intensity × 100%).

### Lentivirus production and transduction

Each sgRNA targeting *TERT* G228A was inserted into a lentiCRISPRv2 vector encoding SpCas9 and sgRNA scaffold (Additional file 1: Fig. S7A). HEK293T cells were prepared on a 10 cm culture dish 24 hours before transfection. Cells were then transfected with 3.75 µg of pMD2.G, 11.25 µg of psPAX2, and 15 µg of lentiCRISPRv2 using 60 µL of Lipofectamine 2000 in 1,500 mL Opti-MEM. After 72 hours, viral supernatants were harvested, filtered through a 0.45-µm PES filter, concentrated, and stored at −80°C. Glioblastoma cell lines were transduced with the lentivirus and subsequently selected with puromycin. After selection, genomic DNA was isolated using the DNeasy Blood & Tissue Kit (Qiagen) for analysis by targeted deep sequencing (Illumina).

#### Targeted deep sequencing

PCR amplified on- and off-target sites for *TERT* G228A using Phusion Hot Start II polymerase (NEB). The primary PCR products were used for a second PCR with NGS adapter primers, followed by a third PCR with indexing primers. PCR amplicons were purified and sequenced by Illumina paired-end sequencing. Sequencing data was analyzed using Cas-Analyzer. Indel frequencies near the PAM site (NGG) were considered CRISPR-Cas9-induced mutations.

#### Single-molecule data acquisition and analysis

For the real-time observation of a dCas9 complex binding to target DNA at the single-molecule level, a custom-built total internal reflection microscopy setup was employed. Cy3-labeled dsRNA was immobilized on a quartz surface coated with polyethylene glycol to minimize nonspecific surface binding. A preassembled complex consisting of dCas9, crRNA and Cy5-labeled tracrRNA was introduced to interact with the immobilized Cy3-labeled dsRNA. A continuous 532 nm laser (Samba100, Cobolt) was used to excite the Cy3 fluorophore on the dsRNA. The fluorescence emission from both the dsRNA and the dCas9 complex was collected using a water-immersion objective lens (60X, N.A.=1.2, Olympus) and then passed through a 542 nm long-pass filter (ET542lp, Chroma) to eliminate the scattered excitation laser light. The filtered fluorescence signal was subsequently split by a 635 nm dichroic mirror (T635lpxr-UF1, Chroma), effectively separating Cy3 and Cy5 emission channels. Each stream of fluorescence was focused onto an EMCCD camera (iXon L-896, Andor) for detection.

The fluorescent intensities of Cy3 (I_Cy3_) and Cy5 (I_Cy5_) were measured from individual fluorescent spots in the single-molecule imaging. The FRET efficiency was calculated using the formula *E* = I_cy5_/(I_cy3_+I_cy5_).

## Declarations

### Ethics approval and consent to participate

Not applicable.

### Consent for publication

Not applicable.

## Supplementary information

Additional file 1 Fig. S1-7

Additional file 2 Table S1-4

