## supplementary figures and tables for "High-fidelity CRISPR genome editing of single-nucleotide mutation with near-complementary guide RNA via enhanced target binding kinetics"

**Additional file 1**

**Figure S1.**

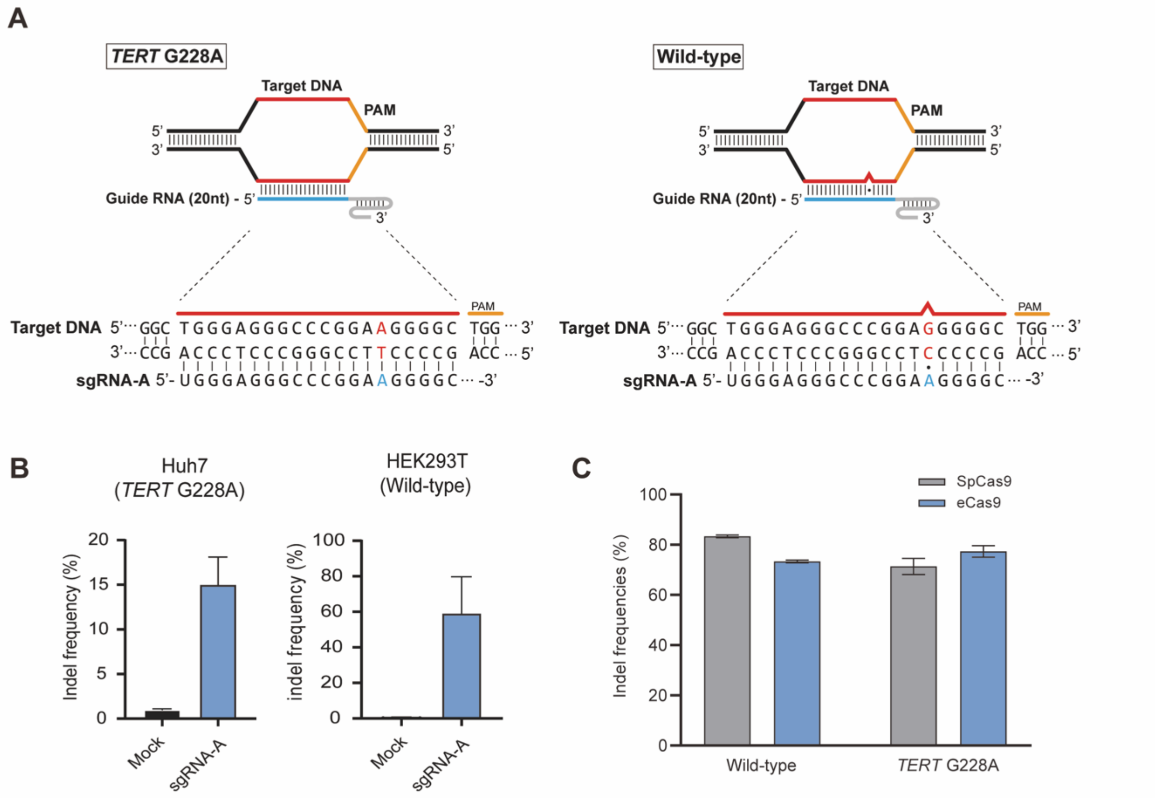


**Figure S1. CRISPR genome editing shows limited ability to discriminate the TERT G228A single-base mutation.** (A) Sequences of the wild-type and TERT G228A mutant alleles. The blue line marks the 20-nt region of the sgRNA complementary to the genomic DNA target sequence (red lines). The enlarged view highlights the TERT G228A point mutation and PAM site, shown above the sgRNA guide sequence. Lines and dots between sgRNA and target DNA bases represent base pairing and mismatches, respectively. (B) Bar graph showing the cleavage activity of the G228A sgRNA against the mutant allele (Huh7) and the wild-type allele (HEK293T). Error bars represent SEM (n = 3). (C) Indel frequencies (%) of SpCas9 and eSpCas9 following co-transfection of sgRNAs with either the wild-type or G228A reporter constructs.

**Figure S2.**

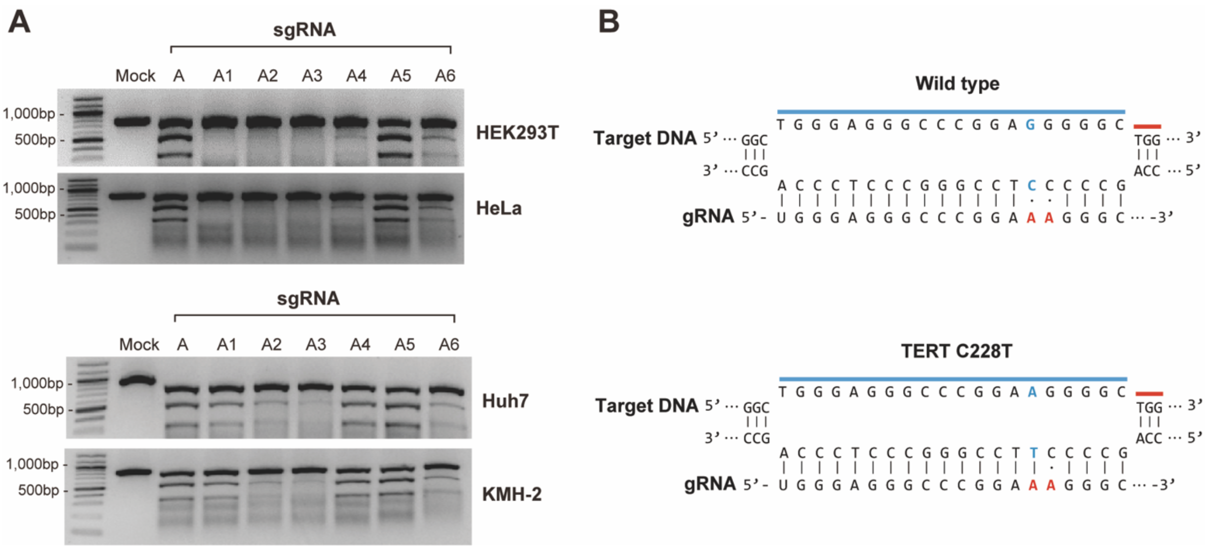


**Figure S2. Identification of near-complementary gRNAs with high specificity to *TERT* G228A mutation by in vitro cleavage assay.** (A) SpCas9 in vitro cleavage assays. Lane (1), Mock (SpCas9 only); lane (2), SpCas9 with sgRNA-A; lanes (3) to (8), SpCas9 with near-complementary gRNA-A1 to -A6. (B) sgRNA-A1 was compared with the wild-type and *TERT* G228A mutant sequences. The red characters show the mismatch sequences to wild type, and the blue characters indicate the TERT G229A mutation region.

**Figure S3.**

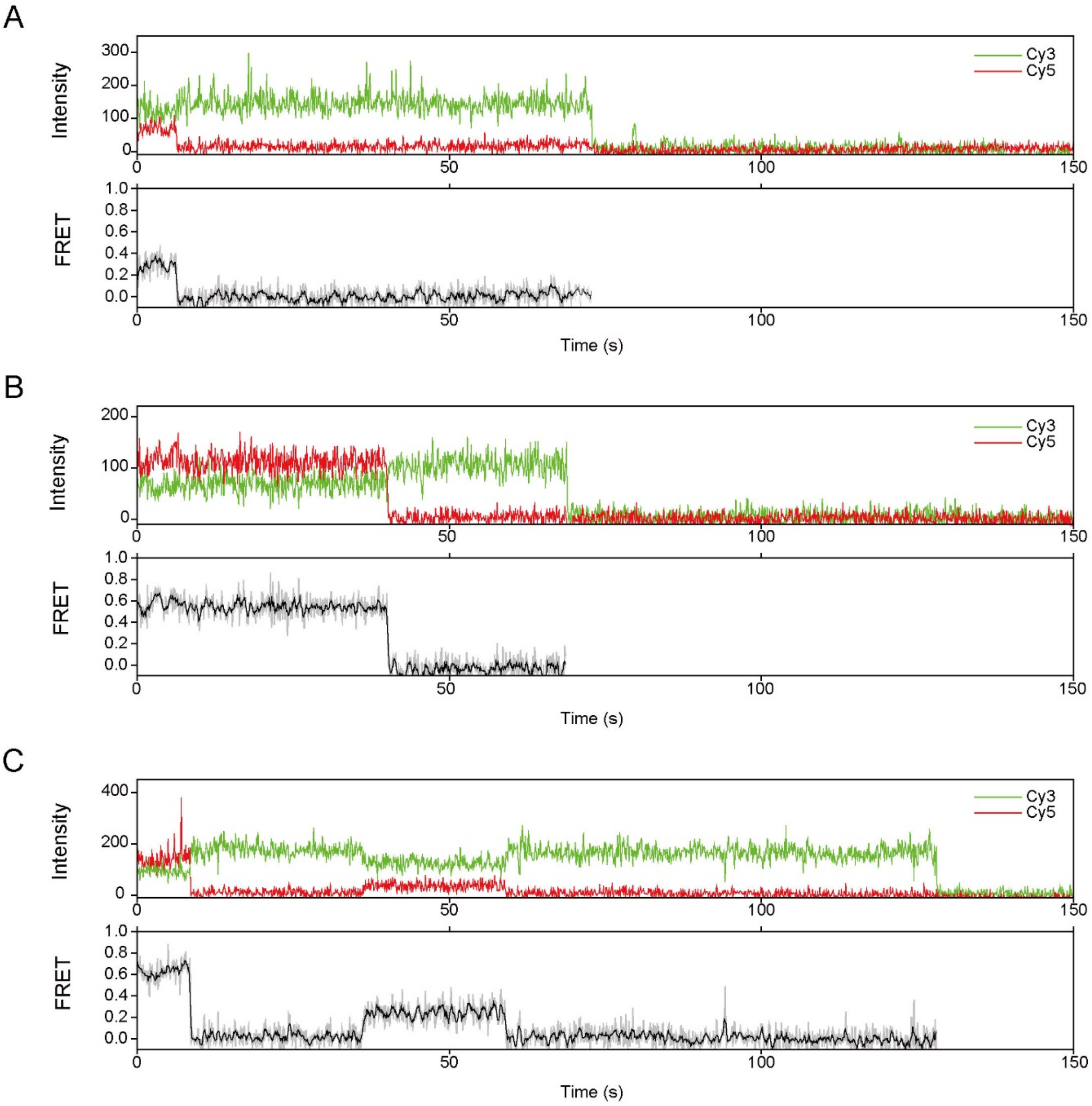


**Figure S3. Single-molecule FRET time traces showing** (A) a steady 0.30 FRET state, (B) a steady 0.55 FRET states, and (C) transitions between the two states.

**Figure S4.**

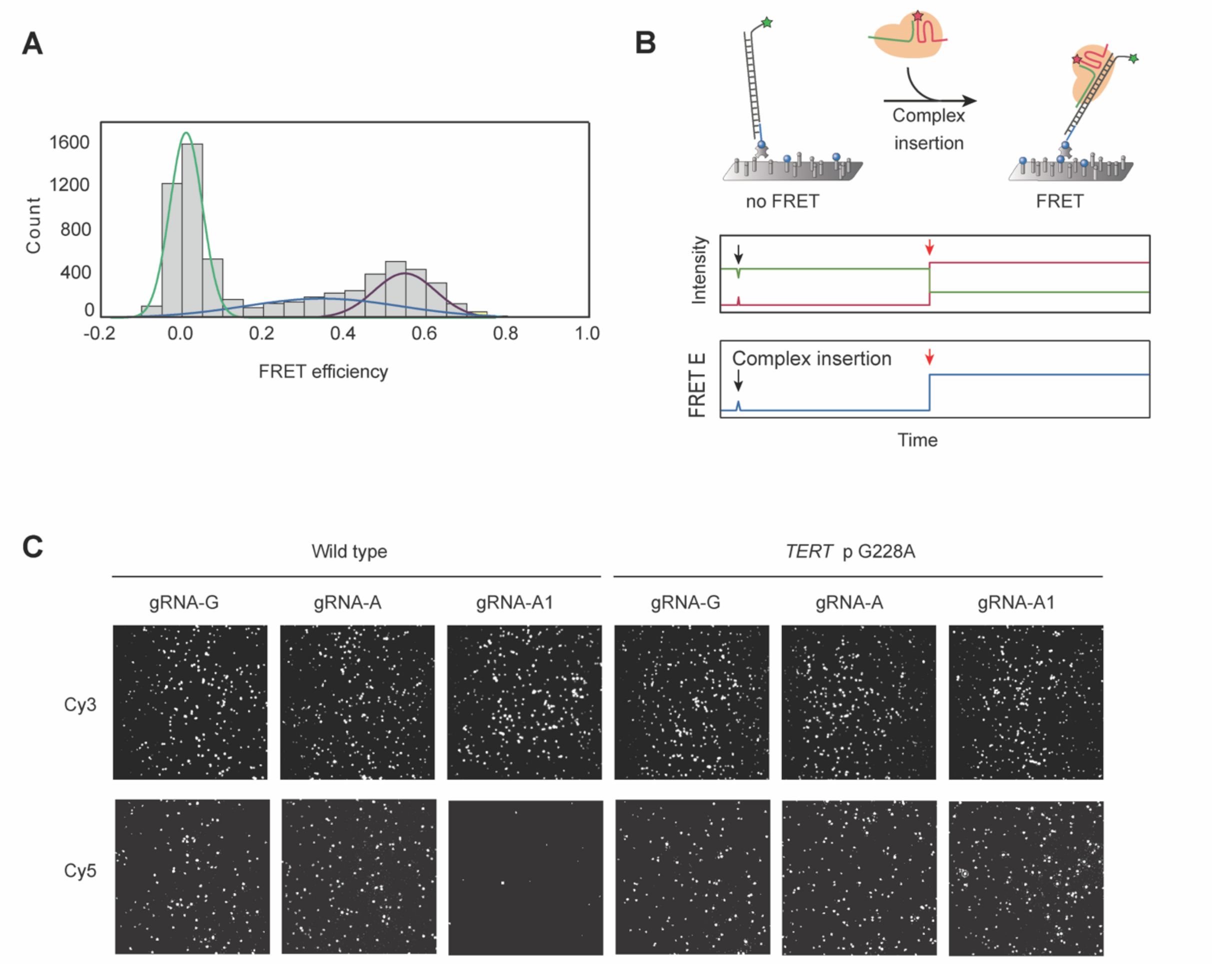


**Figure S4. Single-molecule Binding assay** (A) FRET histogram showing two distinct binding conformations of dCas9-gRNA-DNA complex. (B) Experimental scheme for the flow-based single-molecule assay to monitor real-time binding events through FRET time trajectories. (C) dCas9 complexes with double mismatches between the target DNA and crRNA exhibited significantly reduced binding.

**Figure S5.**

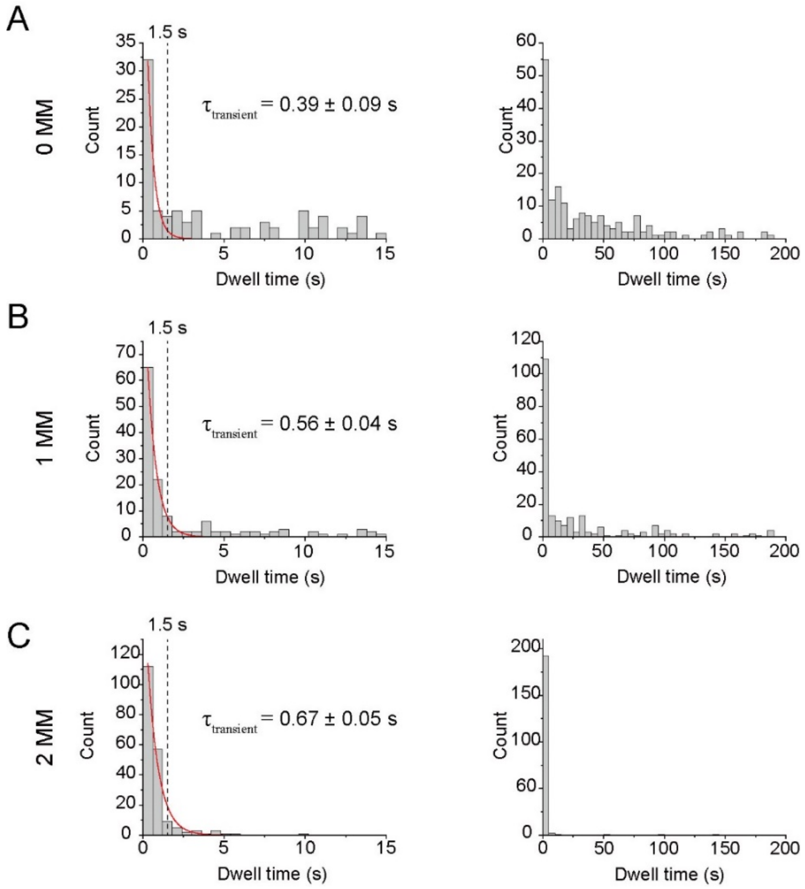


**Figure S5. Dwell time histograms of dCas9 binding to target DNA for (A) no mismatch (0 MM), (B) single mismatch (1 MM), and (C) double mismatches (2 MM) between crRNA and target DNA**. For each condition, the left panels show a partial range of dwell times to highlight the short-lived binding population, while the right panels show the full range of dwell times to display the overall distribution. A short-lived transiting binding population was observed in all conditions, with similar dwell times (0.39, 0.56 and 0.67 s) obtained by fitting to a single-exponential decay (red lines in the left panels). In contrast, the long-lived binding population (>1.5 s) decreased with increasing number of crRNA-DNA mismatches (right panels).

**Figure S6.**

A

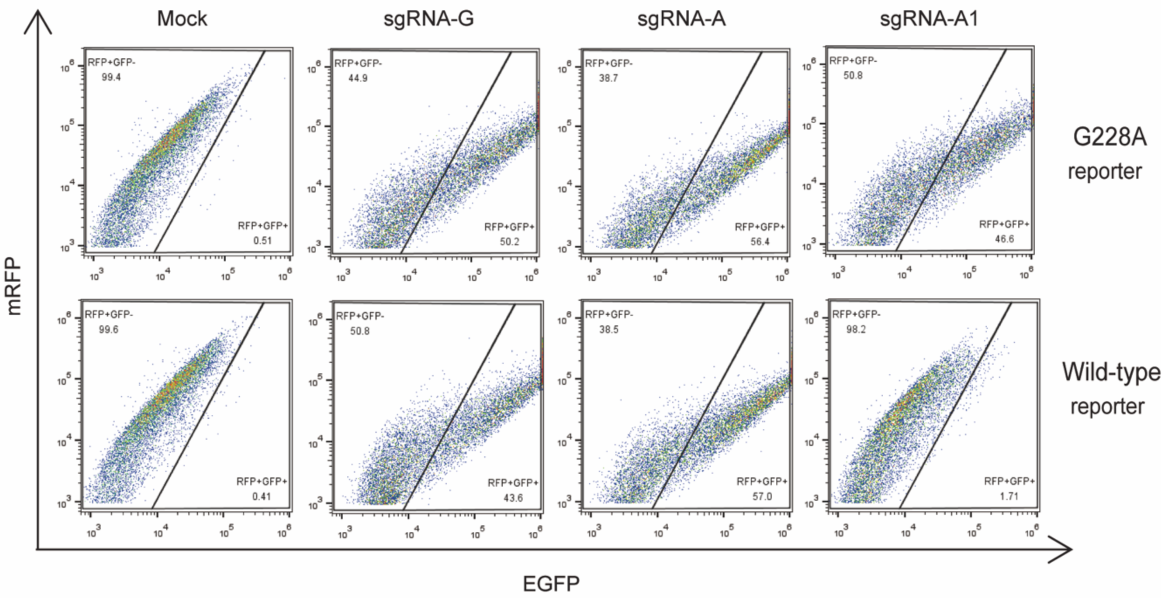


**Figure S6. Dual-fluorescence reporter assay measuring the specificity of near-complementary sgRNAs targeting the TERT G228A mutation.** (A) Flow cytometric analyses to compare the target specificities of sgRNAs targeting the mRFP–EGFP dual fluorescence reporters. The mRFP–EGFP fusion protein was significantly expressed when indels occurred in *TERT* G228A by sgRNA-A1.

**Figure S7.**

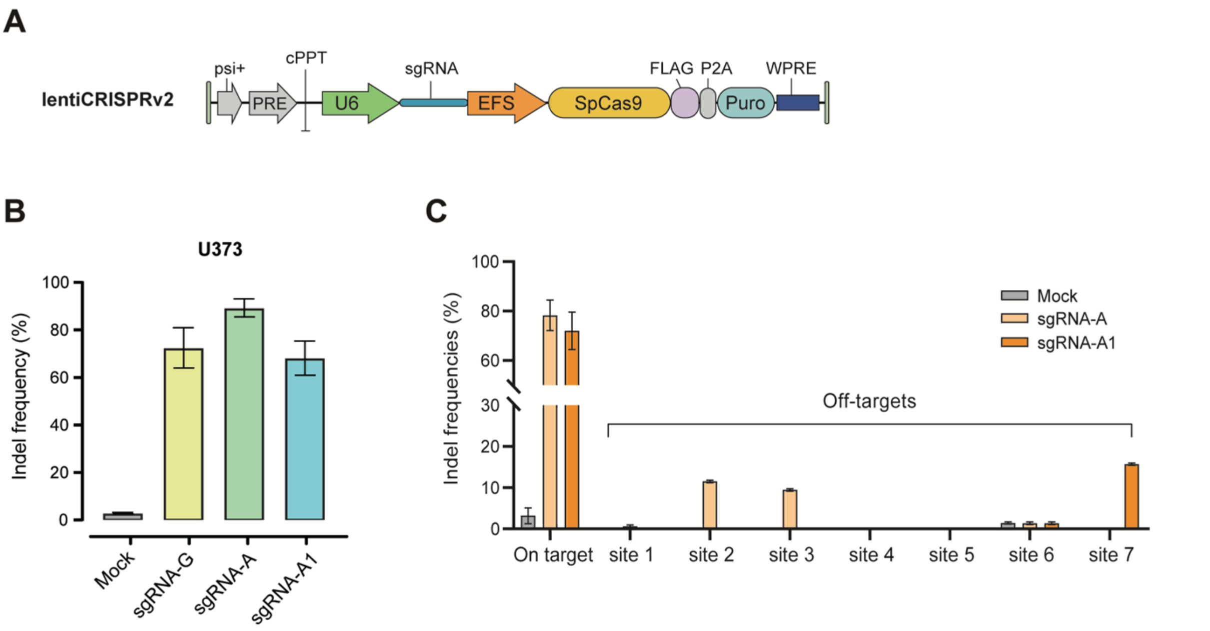


**Figure S7. Single-base precision editing of the endogenous TERT G228A mutation in glioblastoma cells using lentiviral delivery.** (A) Schematic of the lentiviral vector encoding Cas9 and a gRNA targeting the TERT G228A mutation. (B) sgRNA target specificity in the U373 (G228A) cell line. Targeted deep sequencing reveals indel formation at the TERT G228A locus. (C) Indel frequencies at the on-target site and seven candidate off-target sites, quantified by targeted deep sequencing.

**Additional file 2**

**Table S1. near-complementary sgRNA candidates sequence information**

| sgRNA | sequence | Mismatch position relative to PAM |
|---|---|---|
| sgRNA-A | TGGGAGGGCCCGGAAGGGGC | 0 |
| sgRNA-A1 | TGGGAGGGCCCGGAAaGGGC | +1 |
| sgRNA-A2 | TGGGAGGGCCCGGAAGGaGC | +3 |
| sgRNA-A3 | TGGGAGGGCCCGGAAGGGGt | +5 |
| sgRNA-A4 | TGGGAGGGCCCGaAAGGGGC | -2 |
| sgRNA-A5 | TGGGAGGGCCtGGAAGGGGC | -4 |
| sgRNA-A6 | TGGGAGGGtCCGGAAGGGGC | -6 |
| sgRNA-G | TGGGAGGGCCCGGAGGGGGC | - |

**Table S2. off-target counts of sgRNA-A and A1**

| Off-target | sgRNA-A | sgRNA-A1 |
|---|---|---|
| mismatch 1 | 1 | 0 |
| mismatch 2 | 6 | 2 |
| mismatch 3 | 76 | 38 |
| mismatch 4 | 694 | 372 |
| mismatch 5 | 5318 | 3155 |

**Table S3. off-target candidates of sgRNA-A and -A1 targeting TERT G228A**

**sgRNA-A**

| Target | sequence |
|---|---|
| on target | TGGGAGGGCCCGGAAGGGGCTGG |
| off-target 1 | TGGGAGGGCCCGGAgGGGGCTGG |
| off-target 2 | TGGGAGGGCCtGGAAGGGGaAGG |
| off-target 3 | TGGGAGGGCCtGGAtGGGGCAGG |
| off-target 4 | TGaGAGGGCCCGGcAGGGGCTGG |
| off-target 5 | TGGGAGGGCCaGGAtGGGGCAGG |
| off-target 6 | TGGGAGGGCCCGGgcGGGGCAGG |
| off-target 7 | TGGGAGGGCggGGAAGGGGCCGG |

**sgRNA-A1**

| Target | sequence |
|---|---|
| on target | TGGGAGGGCCCGGAAGGGGCTGG |
| off-target 1 | TtGGAGGGCCaGGAAAGGGCTGG |
| off-target 2 | TGGGAGGGCCCGGAggGGGCTGG |

**Table S4. Primers used in experiments**

| primer | sequence(5'-3') |
|---|---|
| TERT _1st_F | GCCGATTCGACCTCTCTC |
| TERT _1st_R | CTCCTTCAGGCAGGACAC |
| TERT _2nd_F | ACACTCTTTCCCTACACGACGCTCTTCCGATCTTGGATTCGCGGGCACAGA |
| TERT _2nd_R | GTGACTGGAGTTCAGACGTGTGCTCTTCCGATCTCAGCGCTGCCTGAAACTCG |
| Reporter _1st_F | TACTTGAAGCTGTCCTTCCC |
| Reporter _1st_R | CCGTCGTCCTTGAAGAAGAT |
| Reporter _2nd_F | ACACTCTTTCCCTACACGACGCTCTTCCGATCTACAACGAGGACTACACCATC |
| Reporter _2nd_R | GTGACTGGAGTTCAGACGTGTGCTCTTCCGATCTCTGAACTTGTGGCCGTTTAC |
| chr15_F | CCATCTGGAGTCATGGAGAAGA |
| chr15_R | GGGAGGAAGGGAGGAAAGTAG |
| chr1_F | CAGTCATGCCCTACAAACGATAAA |
| chr1_R | CTGAGGCATAGTCTCAAGAGGT |
| chr2_F | GGGCTTAGGACTTCGACATAGA |
| chr2_R | GAGTCGCCTTGCTTCCTTTAC |
| chr3_F | GCTCATCGATCACCCACTTAAT |
| chr3_R | CTCGGTCTCCGGGTAACA |
| chr7_F | CTGCCATGCAAGTCCCTATAATC |
| chr7_R | CTCATTCTCCCTCCTCTCATTCA |
| chr9_F | CTCACACCTTCTTGGCTGATAC |
| chr9_R | GGCCGCCTGGAAACTTAAA |
| chr17_F | AGCCTCTGTCCCTGCATAAA |
| chr17_R | CTGACCTGAGGGTCAACATCTTA |
